# GuidingNet: revealing transcriptional cofactor and predicting binding for DNA methyltransferase by network regularization

**DOI:** 10.1101/2020.06.02.129445

**Authors:** Lixin Ren, Caixia Gao, Zhana Duren, Yong Wang

## Abstract

The DNA methyltransferases (DNMTs) (DNMT3A, DNMT3B, and DNMT3L) are primarily responsible for the establishment of genomic locus-specific DNA methylation patterns, which play an important role in gene regulation and animal development. However, this important protein family’s binding mechanism, i.e., how and where the DNMTs bind to genome, is still missing in most tissues and cell lines. This motivates us to explore DNMTs and TF’s cooperation and develop a network regularized logistic regression model, GuidingNet, to predict DNMTs’ genome-wide binding by integrating gene expression, chromatin accessibility, sequence, and protein-protein interaction data. GuidingNet accurately predicted methylation experimental data validated DNMTs’ binding, outperformed single data source based method and sparsity regularized methods, and performed well in within and across tissue prediction for several DNMTs in both human and mouse. Importantly, GuidingNet can reveal transcription co-factors assisting DNMTs for methylation establishment. This provides biological understanding in the DNMTs’ binding specificity in different tissues and demonstrate the advantage of network regularization. In addition, GuidingNet achieves good performance for chromatin regulators’ binding other than DNMTs and serves as a useful method for studying chromatin regulator binding and function. The GuidingNet is freely available at https://github.com/AMSSwanglab/GuidingNet.

**Author summary:** DNA methyltransferases (DNMTs) are in charge of the addition of methyl groups to cytosine residues by binding to DNA specific region. However, DNMTs do not have DNA binding domains to recognize specific DNA sequences and an urging question is how DNMTs recognize their binding sites in the genome in different tissues. Here, we propose a network regularized logistic regression model, GuidingNet, for predicting DNMT’ genome-wide binding by integrating gene expression, chromatin accessibility, sequence, and protein-protein interaction data. The main contribution is to hypothesize that DNMTs interact with transcription factors and are guided by the TF network to bind to DNA, and methylate DNA GuidingNet captures the mechanism of DNMT binding in different tissue contexts and predict DNMTs’ binding well.

## Introduction

DNA methylation is essential for mammalian development and plays crucial roles in various biological processes, including regulation of gene expression, maintenance of genomic stability, genomic imprinting, and X chromosome inactivation [1, 2]. DNA methyltransferases (DNMTs) are in charge of the addition of methyl groups to cytosine residues by binding to DNA specific region. It’s well known that DNMT3A and DNMT3B alone or in a complex with DNMT3L De novo establish DNA methylation, whereas DNMT1 mediates DNA methylation maintenance [3-6].

An urging question is how DNMTs recognize their binding sites in the genome in different tissues. DNA methylation established by these enzymes is highly locus specific and tissue specific but DNMTs do not have binding and tissue specificity by serving as general chromatin regulators. ChIP-seq experiments can measure the genome-wide mapping of DNMTs’ binding locations in specific cellular context [7]. However, ChIP-seq technique requires a large amount of sample material and high quality antibody. Thus DNMTs’ genomic binding in most tissues are still missing. We observed that large scale transcriptomic and epigenomic data across tissues are rapidly accumulated in ENCODE and ROADMAP database. This motivates us to develop computational method to integrate the available omics data, predict DNMTs’ binding, and further understand their binding specificity mechanism.

Our recent study suggests that chromatin regulator (CR) is likely to be recruited to a regulatory element (RE) if RE is open and is bound by transcription factors (TFs), which have protein interaction propensity with the CR (Duren, et al., 2017). This allows us to hypothesize that DNMTs’ binding is guided by TFs to a particular locus to methylate cytosines. TFs is known to express specifically in certain tissue types, recognize specific DNA motifs, bind regions with open chromatin structure, and have direct and indirect protein-protein interactions with DNMTs. In addition, DNA methylation is almost exclusively found in CpG dinucleotides and it suggests DNMTs tend to bind to GC-rich sequences. Taken together, it’s promising to integrate those evidence from available genome sequence, gene expression, chromatin accessibility, protein-protein interaction data to reveal DNMTs’ binding sites and mechanism for binding and tissue specificity.

Collecting the existing DNMTs’ ChIP-seq data as gold standard positives, we model the DNMTs’ binding site prediction as a binary classification problem. The purpose is to mine the matched expression and accessibility data (i.e., measured on the same sample), and context non-specific data (sequence data and protein-protein interaction data), to recover a significant portion of the information in the missing data on DNMTs’ binding location. Numerous statistical methods have been successfully applied in binary classification and logistic regression (LR) is a powerful discriminative method. LR provides predicted probabilities of class membership and easy interpretation of the feature coefficients. In our case, we need to select TFs to provide rich interpretation of the DNMTs’ binding mechanism. Regularized logistic regression (RLR) provides different choices of regularization terms such as the lasso (*ℓ*_1_ – norm) regularization [8] and the elastic net [9], which generalizes the lasso by adding an *ℓ*_2_ penalty. Furthermore, the adaptive lasso regularizes different coefficients in the *l*_1_ - regularization [10] and provides a very general framework for setting feature weights [11-15].

In this paper, we develop a network regularized logistic regression framework, GuidingNet, to predict DNMTs’ binding by integrating multiple dataset. Our major contribution is to reconstruct TF protein-protein interaction and co-expression network for regularization, to choose weights efficiently in adaptive lasso, and to improve prediction accuracy and biological interpretability. GuidingNet is used to predict binding of Dnmt3b and Dnmt3l for E14 mESC, DNMT3A for human MCF-7 cells, and DNMT3B for human HepG2 cells and foreskin keratinocyte. GuidingNet has a superior performance in both the prediction and feature selection. Also GuidingNet outputs a TF co-factor network to interpret the mechanism of DNMTs’ binding specificity in genomic locus and in different tissue contexts.

## Materials and methods

### Overview of GuidingNet model

We propose GuidingNet to model the physical process that TF recruits DNMTs to a specific chromatin region to methylate cytosines (S1 Fig), i.e., DNMT is guided by TF to recognize the specific DNA motifs and bind in a RE. This requires that RE should be accessible to TF binding and TFs have protein interaction propensity with the DNMT. GuidingNet takes the context specific and non-specific genomic data as input and outputs the DNMTs’s binding probability on a given regulatory element and the guiding TF network. As depicted in Fig 1, GuidingNet has three components respectively, (i) extracting predictive genomic features and combining feature with physical meaning, (ii) generating the candidate guiding TF network, and (iii) training the model and outputting DNMT binding and TF cofactor network.

**Fig 1.**
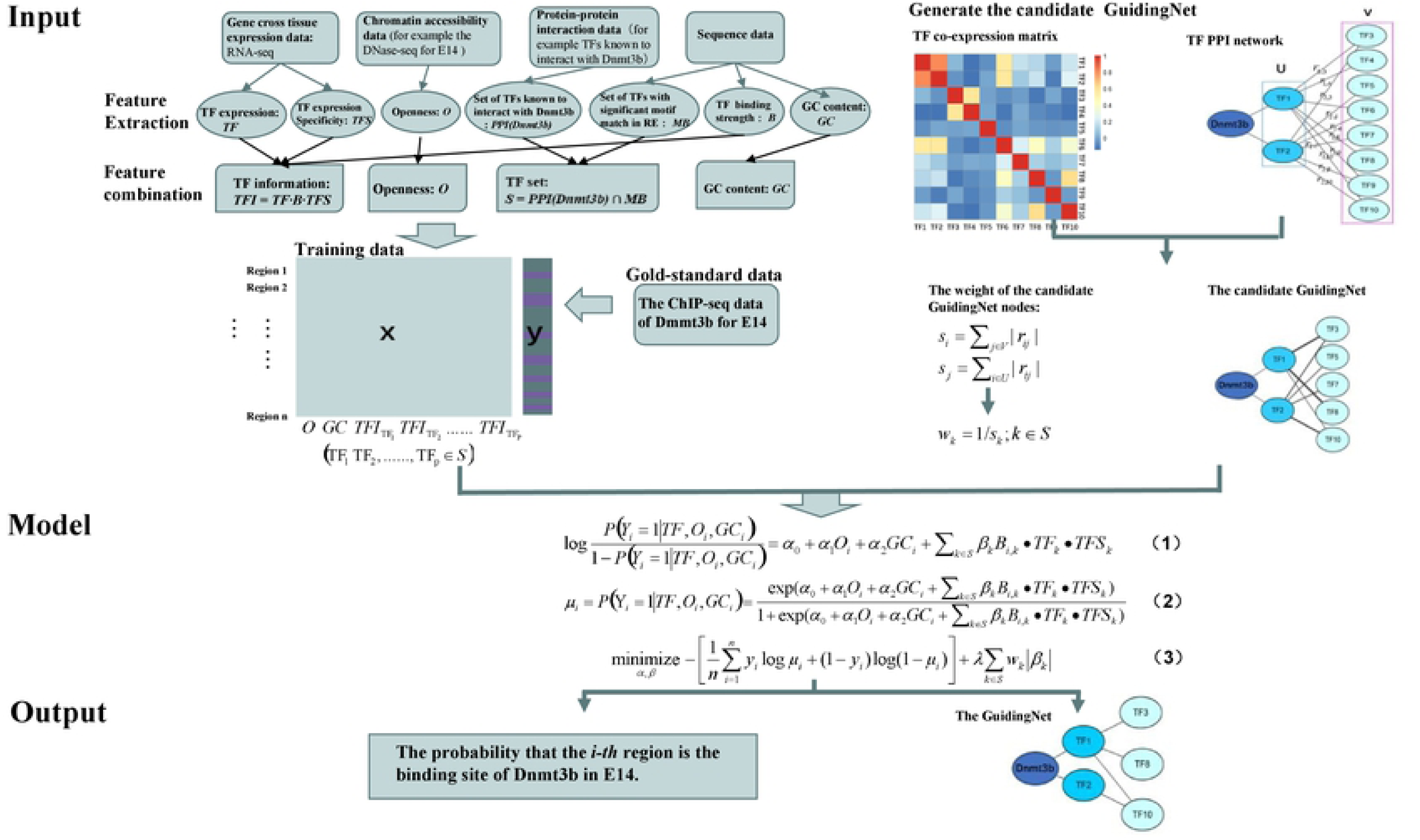
Overview of GuidingNet model. GuidingNet is illustrated from three parts: input, model, and output. Dnmt3b in mouse E14 is used an example for DNMTs’ binding prediction. Model input includes the construction of training data and the candidate GuidingNet. Openness, expression, sequence, and protein-protein interaction data are collected and processed as the GuidingNet’s input. Seven genomic features are extracted from the input data. TF expression, TF expression specificity, and TF binding strength are further combined as one feature. This together gives training data X. The training labels are from the ChIP-seq data. Meanwhile the candidate GuidingNet is generated and weighted base on the TFs’ protein-protein interaction network and co-expression. GuidingNet is trained by an adaptive lasso-regularized logistic regression framework with the candidate GuidingNet and corresponding training data. The model components are described in Table 1 with mathematical notations. Model output are the probability that the region *i* is the binding site of Dnmt3b in E14 and the underlying GuidingNet.

Fig 1 shows the example for Dnmt3b’s binding prediction in mouse E14. The input data include chromatin openness, expression, sequence, and protein-protein interaction. Features are extracted from the input data to construct training dataset. The training labels are from ChIP-seq data. Independently, the candidate GuidingNet are generated and weighted based on the TF protein-protein interaction network and co-expression. We used the TFs and weights of the candidate GuidingNet to train a network regularized logistic regression model. Model output are the probability that a specific region is the binding site of Dnmt3b in E14 and the underlying GuidingNet. The model components are described in Table 1 with notations.

**Table 1.** Model components for GuidingNet.

| Description of data and variables | Notation | Example |
| --- | --- | --- |
| Expression of TF | $TF_k$: expression of TF $k$ | $TF_{Nanog} = 30.95$ in E14 mESC |
| Accessibility of a genomic region | $O_i$: degree of openness of region $i$ | $O_{chr7:12,922,178 - 12,922,505} = 12.83$ in E14 mESC |
| Binding status of Dnmt3b in a genomic region | $Y_i$: indicator for whether the Dnmt3 is binding to region $i$. | Dnmt3b binds genome at chr7:12,922,178-12,922,505 in E14 mESC |
| Interacting TFs for Dnmt3b | $PPI(Dnmt3b)$: sets if TFs known to interact with Dnmt3b | $PPI(Dnmt3b)$ contains Nanog |
| TFs with motif match in a genomic region | $MB$: set of TFs with significant motif match in region $i$ | Nanog has motif match at chr7:12,922,178-12,922,505 in E14 mESC |
| Motif matching strength of TF in a genomic region | $B_{i,k}$: binding strength of TF $k$ in region $i$ | $B_{chr7:12,922,178 - 12,922,505, Nanog} = 5.39$ |
| GC fraction of a genomic region | $GC_i$: GC content of region $i$ | $GC_{chr7:12,922,178 - 12,922,505} = 0.61$ |

### Feature collection and combination

We extracted five features including openness (*O*), TF binding strength (*B*), TF expression (*TF*), TF expression specificity (*TFS*), and GC content (*GC*) from our input data (see details in Supplementary Materials). These features are not independent. Different combinations indicate differential recruitment pattern of DNMT binding to the RE. We used the following feature transformation to model the three physical steps for the DNMTs binding processes that: 1) region is open, 2) TF binds to the region, and 3) region has proper GC content for methylation. *TF•B•TFS* indicates that TF has significant motif match on the RE, highly expressed and specific expression across tissues (Fig 1). Then we have three features openness (*O*), TF binding information (*B•TF•TFS*), and GC content (*GC*) for each regulatory region.

### Candidate GuidingNet generation

The generation of candidate GuidingNet consists of two steps (Fig 1 for Dnmt3b in mouse E14 example):

**Step 1**. Define a bipartite graph *G*(*U, V, E*) based on Dnmt3b PPI network structure. TF set is constructed as *S* = *PPI* (*DNMT* 3) ∩ *MB* by considering protein-protein interaction (PPI) with DNMTs and motif occurrence. S is further divided into U and V. The set of TFs having direct PPIs (first order neighbor) with Dnmt3b defined as *U*. The set of TFs having indirect PPIs (second order neighbor) with Dnmt3b defined as *V*. The edges of the graph are defined if TF *i* in U has PPI with TF *j* in V. We use *c*_*ij*_ to represent the weight of the edge. *C*_*ij*_ is calculated by the correlation between TF *i* and TF *j* for the expression levels across diverse tissues (as shown in Fig 1).

**Step 2**. Set cutoff *Q*,

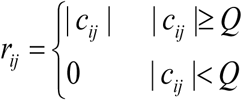

Let *s*_*i*_ = Σ_*j*∈*V*_ | *r* _*ij*_ |, and *s*_*j*_ = Σ_*i*∈*U*_ | *r*_*ij*_ |. We then construct feature weights *w*_*k*_ = 1/ *s*_*k*_ ; *k* ∈ *S* and generated the candidate GuidingNet.

### Network regularized logistic regression

We model the DNMTs’ binding to regions of the genome by a logistic regression, which is a statistical method for a binary classification problem. Given a DNMT, we can measure its ChIP-seq data and extract peaks as the gold standard positive data (GSP) for binding. The gold standard negative data (GSN) are randomly sampled from the non-binding regions (see details in Supplementary Materials). Assume the entire training gold-standard data (includes GSP and GSN) and genomics feature data have n regions and p predictors. We denote whether a DNMT3 binding to regions *i* as *Y*_*i*_ ∈{0,1}, *Y*_*i*_=1 means region *i* is bound by DNMT and *Y*_*i*_=0 means not. The genomic features are openness (*O*), TF binding information (*B• TF• TFS*), and GC content (GC).

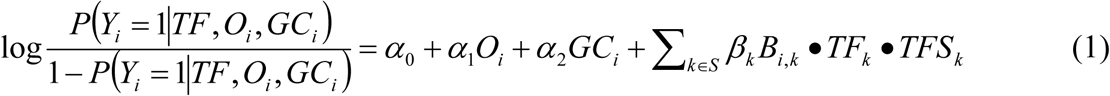

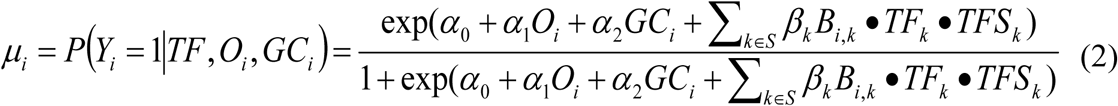

Where *S* = *PPI* (*DNMT* 3) ∩ *MB*. The log-likelihood function of Eq. (1) is defined as:

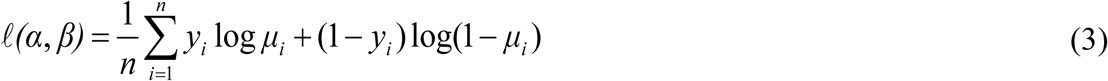

where *α*= (*α*_0_, *α*_1_, *α*_2_) is a vector of TF independent coefficients, *β*= (*β*_1_, *β*_2_, ……) is a vector of TF dependent coefficients, and *y*_*i*_ ∈{0,1} is the training label of region i. We can estimate the parameters by minimizing the negative log-likelihood function.

With the candidate GuidingNet, Our network regularized logistic regression optimizes the adaptive lasso of *β* by:

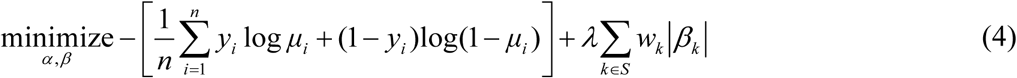

where *w*_*k*_ is the weight of the TF information of *k*. In our case, the weights of predictors are related to TFs. However, these TFs are not independent and they may have physical interactions. To handle this, we purpose GuidingNet by choosing weights based on the TF protein-protein interaction network and co-expression in an unsupervised manner. In practice, we fit the network regularized model using the vector of weight as “penalty factors” in the glmnet R package and generated the GuidingNet and probability that a given region is the binding site of DNMT.

### Data sources

We collected the ChIP-seq of DNA methyltransferases 3 (DNMT3) family proteins including Dnmt3b and Dnmt3l for E14 mouse embryonic stem cells (mESC), DNMT3A for human MCF-7 cells, DNMT3B for human HepG2 cells and foreskin keratinocyte form the Cistrome Data Browser (http://cistrome.org/db). The corresponding paired RNA-seq and DNase-seq are downloaded from the ENCODE project. We collected RNA-seq data from other 145 mouse (S1 Table) and 832 human (S2 Table) samples from the ENCODE and ROADMAP. We also collected the WGBS data of mouse E14 [16]. Both mouse and human protein-protein interaction data are from the BIOGRID database (https://thebiogrid.org). The RNA-seq, DNase-seq and WGBS of mouse embryo development are downloaded from the ENCODE project.

## RESULT

### Openness, TF information, and GC content are predictive for DNMTs’ binding

We first quantitatively assess the usefulness of openness, TF information and GC content feature for DNMT3 binding prediction by area under the curve (AUC) value with univariate ordinary logistic regression. As shown in Fig 2, all three genomic features are good predictors for the DNMT3 prediction in five scenarios. All AUC values of single feature predicting exceed 0.5. The performance of openness, TF information, and GC content of binding prediction for the same DNMT3 protein in different cell types and different DNMT3 proteins were distinctive. In particular, the predictive ability of TF information is great except in human HepG2, demonstrating that the importance of TF guidance in the process of DNMT3 binding to the regulatory elements. The predictive ability of openness and GC content are varied in different cell types. For example, the AUC for openness prediction in DNMT3A for MCF-7 is 0.82, but only 0.53 in Dnmt3b for mouse E14. The AUC for GC content in Dnmt3b for E14 is 0.85, but only 0.59 in Dnmt3l for mouse E14. For DNMT3B, three feature performance are different in E14 mESC, human foreskin keratinocyte, and HepG2 cells. This illustrates that the same DNA Methyltransferase had a distinctive binding pattern in different cell types.

**Fig 2.**
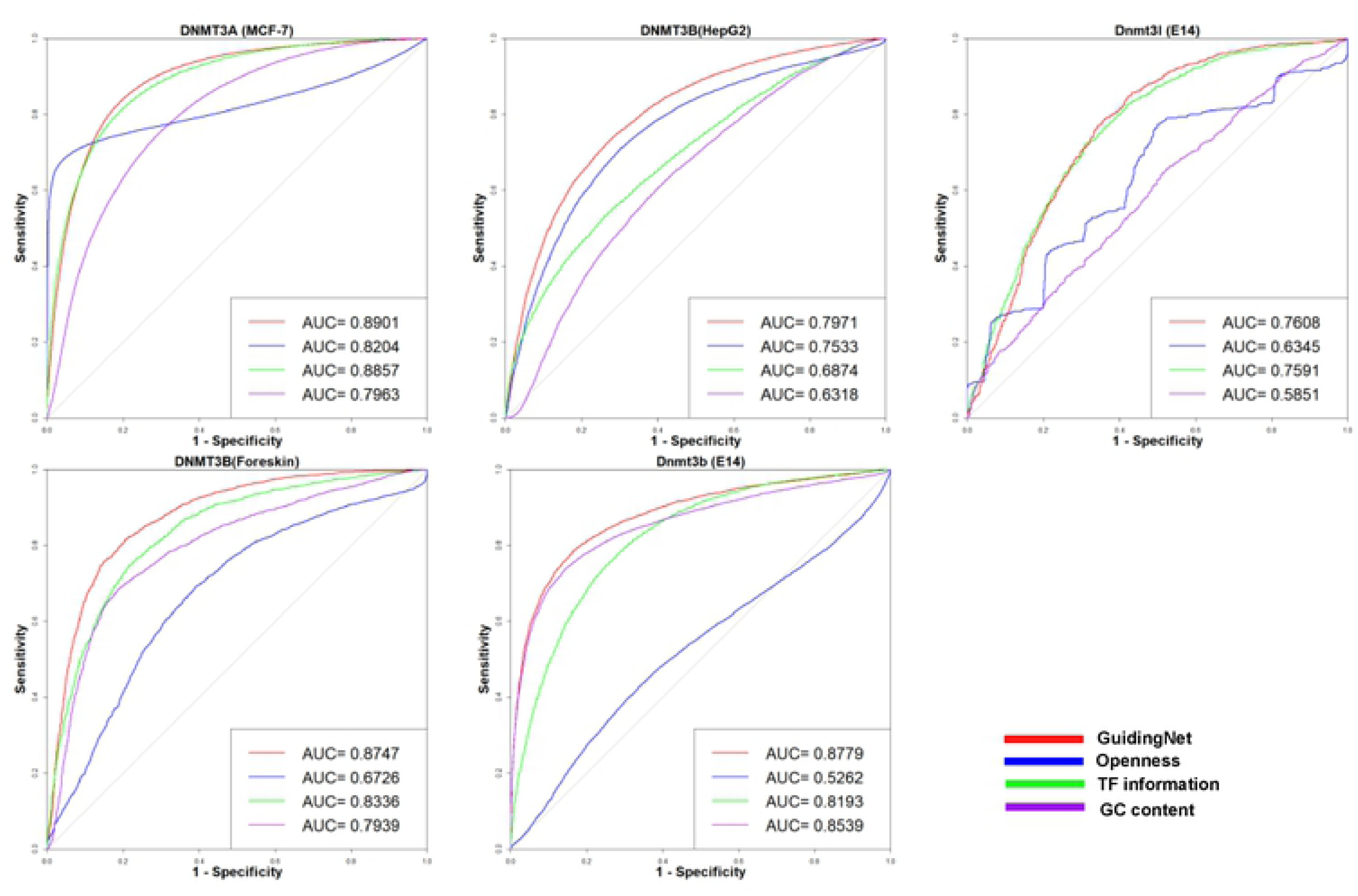
Every single feature has predictive power and GuidingNet is better than that base on single feature for predicting the DNMT3 family proteins binding. ROC curves of single feature and GuidingNet prediction of binding for DNMT3A in human MCF-7 cells, DNMT3B in human HepG2 cells, DNMT3B in human foreskin keratinocyte, and Dnmt3b and Dnmt3l in E14 mESC.

### GuidingNet improves binding prediction

Our GuidingNet holds the promise to provide biological insights in the DNMTs’ binding mechanism in different cellular contexts. We systematically evaluated the performance of in prediction and feature selection for the above five scenarios. As indicated in Fig 2, ROC curves of the GuidingNet prediction on these five scenarios show better performance, i.e., averagely 84 % area under the curve. Importantly, GuidingNet is better than the predictions based on unitary feature. This demonstrated that the importance of integrative multi-omics data for predicting DNA methyltransferase binding.

### GuidingNet reveals biological meaningful TF cofactors to enhance interpretability

The GuidingNet is a two-step way for feature selection and shows good performance. As shown in Fig 3A-D, the candidate GuidingNet includes fewer TFs than protein-protein interaction network and the cutoff set was 0.1 for mouse and 0.7 for human cells. For example, the candidate GuidingNet of DNMT3A for the MCF-7 cells included approximately one quarter of TFs in the protein-protein interaction network. Then we further filtered the candidate GuidingNet when fitting the network regularized logistic regression. Our method selected 20.8%, 15.6%, 16.8%, 53%, and 16.3% TFs of protein-protein interaction network in DNMT3A for human MCF-7 cells, DNMT3B for human HepG2 cells and foreskin keratinocyte, Dnmt3b and Dnmt3l for E14, respectively (Fig 3A-D). Comparing with ordinary logistic regression (OLR) using openness, GC content, and PPI network in Fig 3A-D (S2 Fig). GuidingNet’s accuracy remains but used far fewer TFs for prediction. For example, our method used one-fifth TFs of OLR for DNMT3B binding in the human HepG2 cells.

**Fig 3.**
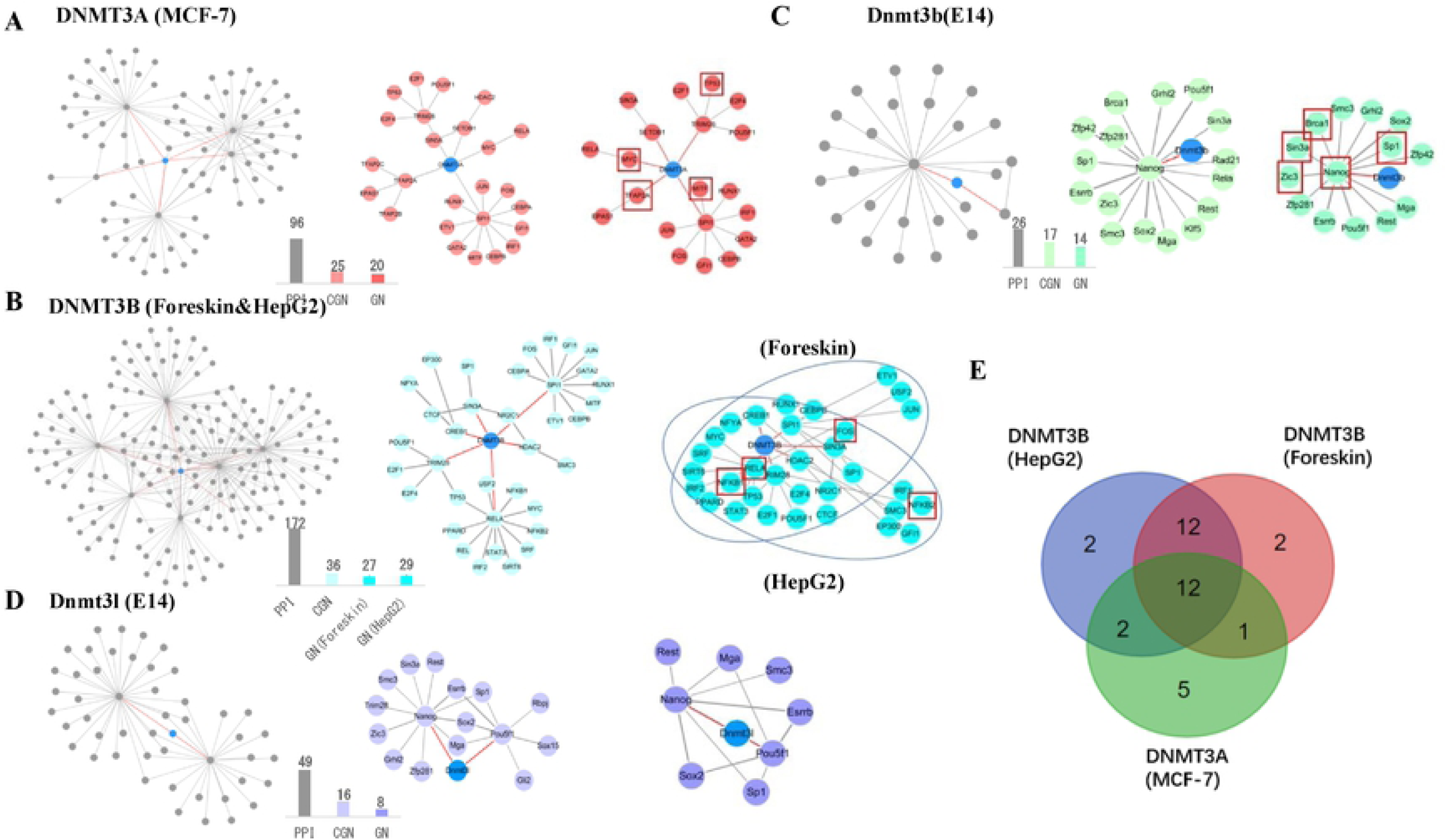
GuidingNet’s performance in selecting less TF co-factors and enhancing interpretability. (A-D) Comparison of protein-protein interaction networks (included first-order and second-order TF interact with the DNMT3 family protein), the candidate GuidingNet, and the GuidingNet of DNMT3A in human MCF-7 cells, DNMT3B in human HepG2 cells, DNMT3B in human foreskin keratinocyte, Dnmt3b in E14 mESC, and Dnmt3l in E14 mESC. The cutoff used to generate the candidate GuidingNet was 0.1 for mouse and 0.7 for human. The blue nodes in protein-protein interaction networks indicate the DNMT. The red edges indicate the direct interaction between the DNMT with TFs. The width of the edge in the candidate GuidingNet indicates its weight. The wider the width of the edge, the greater its weight. Bar plot indicates the number of TFs in the PPI network (PPI), the candidate GuidingNet (CGN), and the GuidingNet (GN). The red box indicates TFs having motif enriched in the DNMT’s ChIP-Sep peaks. (E) Venn diagram shows the number of shared TFs among the GuidingNet of DNMT3A in MCF-7, DNMT3B in HepG2, and DNMT3B in foreskin keratinocyte.

The co-factors in the output GuidingNet we called the guiding co-factors and can help us interpret the mechanism of how co-factor assist DNA methyltransferase binding to regulatory elements. To validate those predicted cofactors, we preformed motif enrichment in the ChIP-seq of DNMT3 proteins by HOMER software. As shown in Fig 3A-D, we indeed identified guiding TFs enriched in the ChIP-seq of corresponding DNMT3. For example, Nanog, Sp1, Brca1, Sin3a, and Zic3 in Dnmt3b for E14. In addition, we searched the literature to show a number of studies have linked our predicted guiding co-factors to DNA methyltransferase. For example, TRIM28-mediated recruitment of de novo DNMT3A leads to cytosine methylation at CpG dinucleotides can control human endogenous retroviruses [17]. A total of 24 guiding co-factors were validated to be associated with DNA methyltransferase binding by literature (S3 Table). We note that the guiding co-factors may play an important role in development and many diseases. Uncovering the mechanisms by which these co-factors regulating DNA methyltransferase binding may be useful in the production of therapeutic targets.

Then we compared the GuidingNets of these DNMT3 family proteins across different cellular context. It is interesting that the GuidingNets of DNMT3A and DNMT3B in human cells shared most of the guiding TFs (Fig 3E). Previous studies have shown that DNMT3A and DNMT3B are structurally similar and appear to have redundant functions overall [18]. One possible reason is that DNMT3A and DNMT3B are recruited by the same TFs. In summary, the GuidingNet shows superior performance in predicting DNMT3 family proteins in both mouse and human and help us to understand the binding co-factors to the REs.

### GuidingNet’s genome-wide prediction is validated by WGBS data

We used the whole genome bisulfite sequencing (WGBS) data on mESC E14 to validate GuidingNet’s Dnmt3b binding prediction. Given a certain region, we can quantify the methylation level for this region by a simple fold change score, which is calculated as the number of methylated cytosine in this region by comparing with the total number of cytosine in this region. As indicated in Fig 4A, the methylation level of Dnmt3b binding regions is higher than not bound Dnmt3b regions in E14 mESC. Fig 4B shows that the higher the methylation level of this region, the higher the probability that this region is a Dnmt3b binding site. In our prediction of Dnmt3b binding, the Pearson correlation coefficient between the binding probability of Dnmt3b to the region and the methylation level of this region is 0.816, indicating that our prediction results are consistent with the independent measured WGBS data.

**Fig 4.**
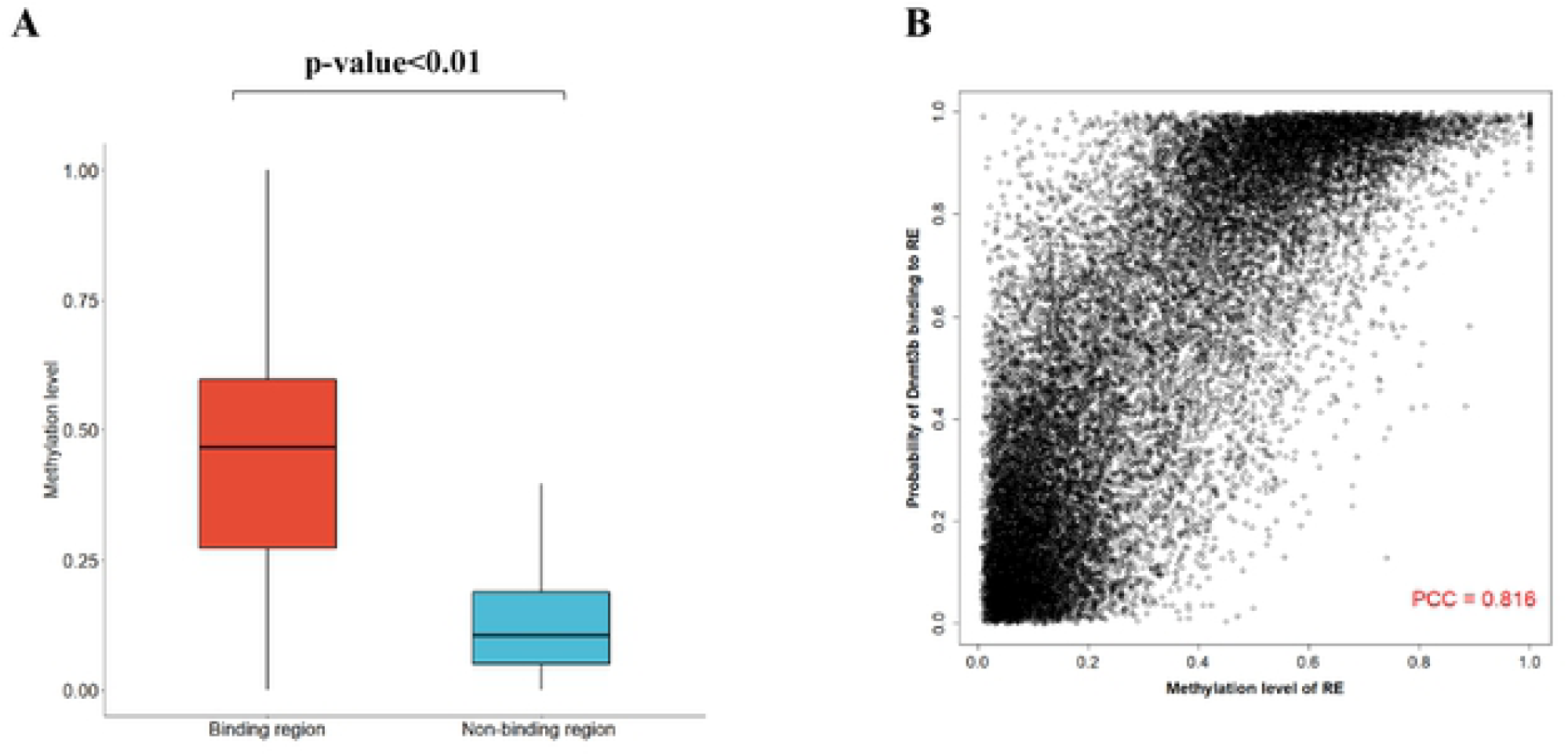
DNA Methylation data validate the Dnmt3b binding prediction in E14 mESC. (A) Methylation level of Dnmt3b’s predicted binding regions is significantly higher than non-binding regions (define as the open region not covered Dnmt3b’s binding region) in E14 mESC. (B) The scatter plot of the probability of adaptive logistic regression for the Dnmt3b’s binding versus the methylation level for a given region.

### Across-tissue predicting Dnmt3b’s binding

To further evaluate our model performance, we performed across-tissue prediction of Dnmt3b binding by GuidingNet. As shown in Fig 5A, we trained the model in E14 mESC then predicted which open regions are Dnmt3b binding sites at 7 time points (E11.5, E12.5, E13.5, E14.5, E15.5, E16.5, and P0) of the liver during embryo development. We used the WGBS data to validate the Dnmt3b’s binding prediction. As indicated in Fig 5B, the methylation level of Dnmt3b’s predicted binding regions is higher than non-binding regions. This is consistent with the results of the WGBS data validation in E14 mESC. These results suggest that our method can apply in across-tissue prediction of Dnmt3b binding. Interestingly, the methylation level of the predicted binding region in E14.5 is higher than other time points. We ask the question that whether the better prediction of E14.5 is related to the right guiding TFs choice. We compared predicted binding regions between E14.5 and E13.5, then extracted the 11,494 unique binding regions of E14.5 (S16A Fig). We did TF motif enrichment in these regions by HOMER and identified 8 motifs for enriched guiding TFs (Nanog, Sp1, Smc3, Zic3, Zfp281, Esrrb, Pou5f1, Rest). More importantly, Smc3, Sp1, Zfp281, and Rest have high expression levels in E14.5 (S16B Fig). In addition, the distance between predicted binding regions and the transcription start site (TSS) at E14.5 is shorter than other time points (S17 Fig). This may indicate that our method trained in mouse E14 is more accurate in the promoter regions. We expect to train our model in more tissue and cell types, and consequently we could predict the DNMT binding for a new tissue and cell type that has not been studied yet.

**Fig 5.**
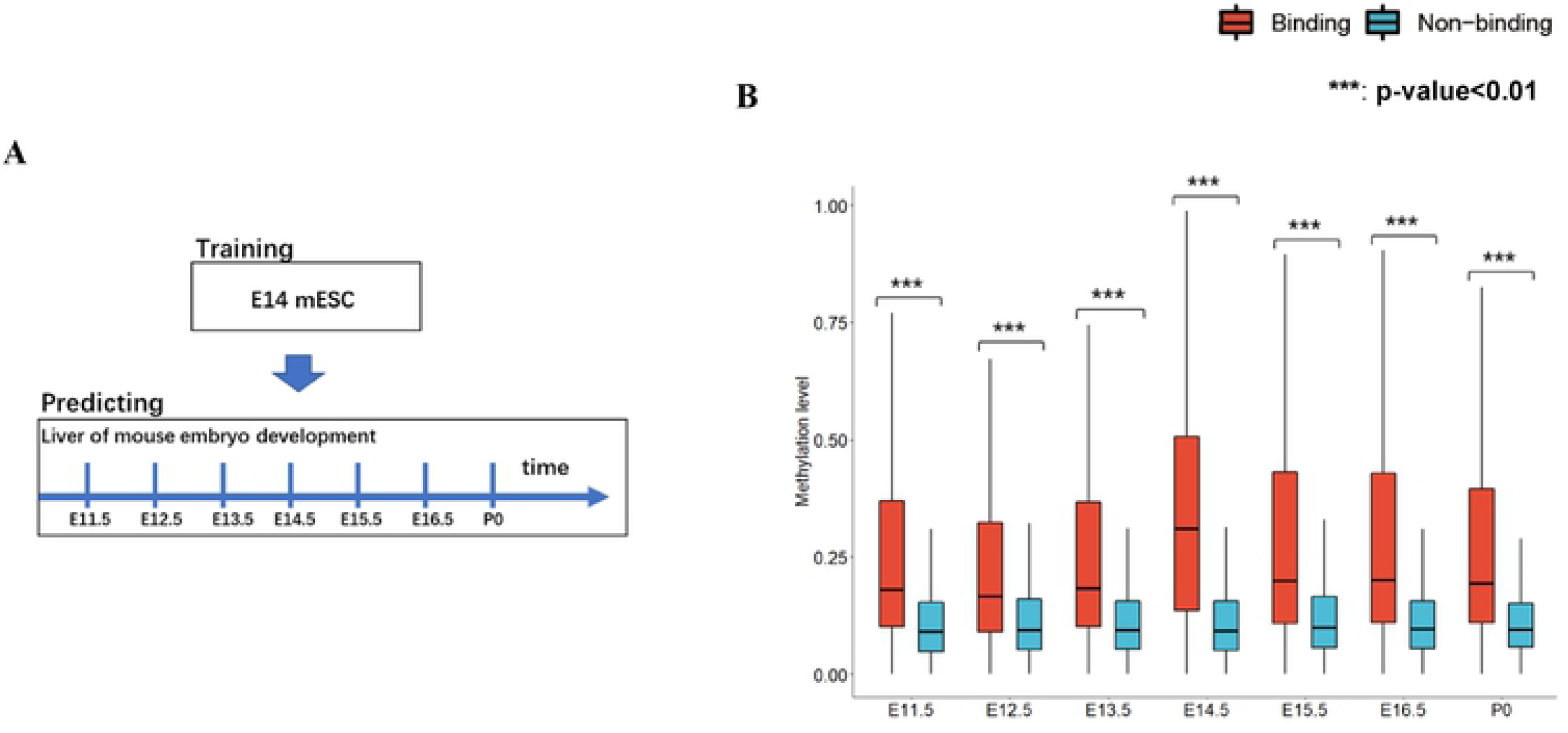
GuidingNet predicts Dnmt3b’s across-tissue binding. (A) Schematic of training the model and across-tissue predicting. (B) Comparison of methylation level between the Dnmt3b’s predicted binding regions and non-binding regions.

### GuidingNet can predict other chromatin regulators’ binding in human and mouse

We ask whether our model has the predictive power in other chromatin regulators’ binding. The ChIP-seq datasets for 4 CRs (Ep300, Ezh2, Setdb1, and Suz12) in mouse E14 and 6 CRs (EP300, EZH2, HDAC2, SMARCC2, SMARCE1, and SUZ12) in human HepG2 cells are available. We evaluated our predictions of the binding of these CRs. Fig 4A and 4B shows the ROC curves of these CRs with good performances (74– 94% AUC) have been achieved. We also assessed the predictive power of openness, TF information, and GC content (S4 Fig and S9 Fig). TF information showed the superior performance for all these CRs’ binding predictions and GC content had weak predictive power. The predictive ability of openness varies for different CRs. In addition, our method showed outstanding performance in TF related feature selection (S5-8 Fig and S10-15 Fig). In particular, our method had shown superior accuracy in predicting SUZ12’s binding in both mouse and human. The AUCs are 0.91 and 0.94 respectively. In addition, a total of 7 TFs (BRCA1, YY1, STAT3, NFATC1, SMC3, POU2F1, and NANOG) are shared in the GuidingNet of human HepG2 (Fig 6C) and mouse E14 (Fig 6D).These TFs are conserved from mouse to human, suggesting that they play an important role in guiding SUZ12 binding to the regulatory elements. The revealed GuidingNet of these CRs in E14 mESC and HepG2 cells are biologically reasonable (S5-8 Fig and S10-15 Fig). The strong performance both in prediction accuracy and TF selection suggests that GuidingNet is useful in other chromatin regulators’ study.

**Fig 6.**
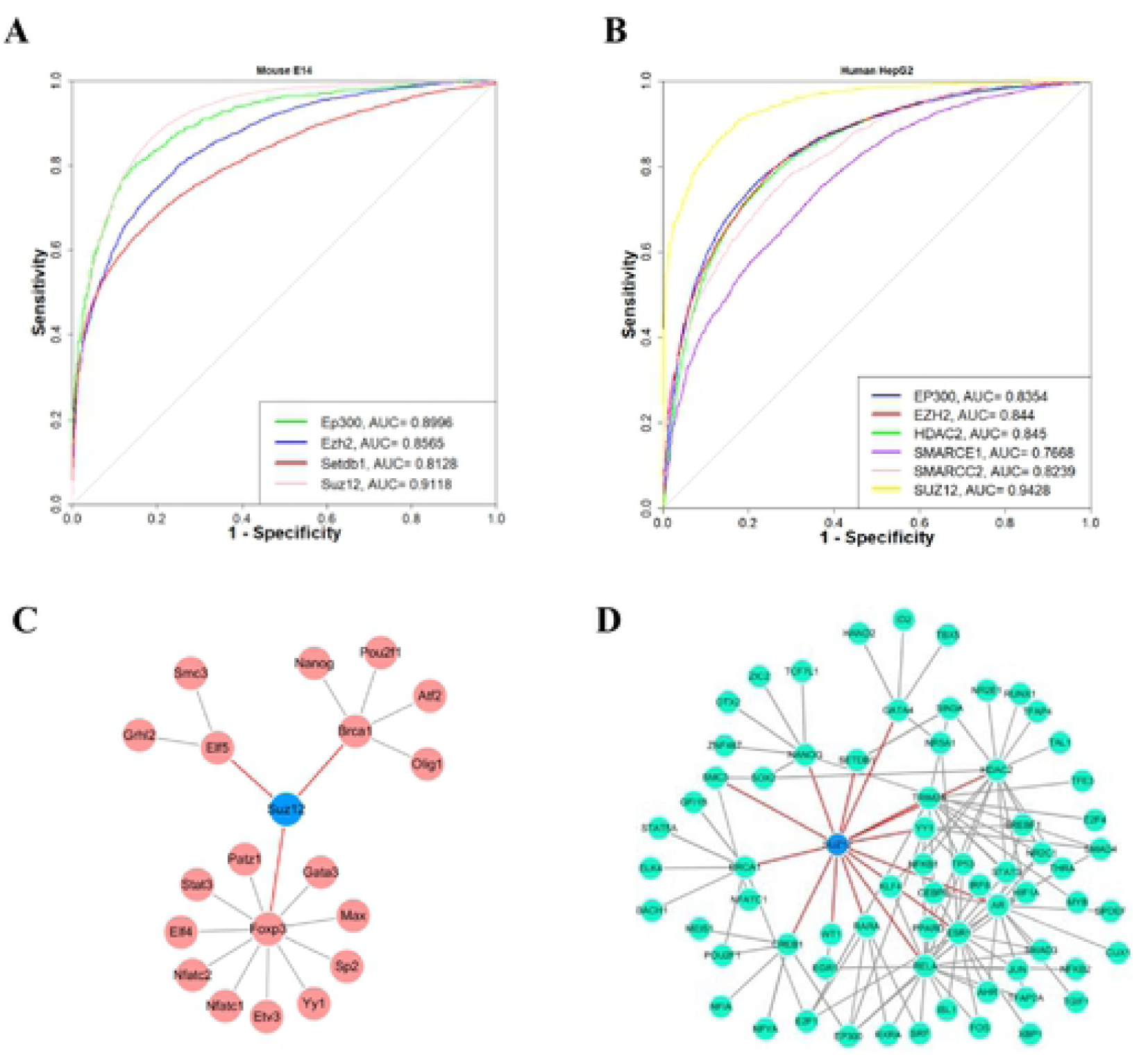
GuidingNet can predict other CR’s binding in human and mouse. (A) ROC curves of prediction of 4 CRs’ binding in the mouse E14. (B) ROC curves of prediction of 6 CRs’ binding in the human HepG2 (C)-(D) The GuidingNet of Suz12 in mouse E14 mESC and SUZ12 in human HepG2, respectively.

## Discussion

We introduced GuidingNet, a network-regularized logistic regression framework, to integrate gene expression data, chromatin accessibility data, DNA sequence information, and protein-protein interaction data for modeling DNMTs’ binding. Our major contribution is to propose a new method to choose TF related feature weights in the adaptive lasso base on TF protein-protein network and co-expression. Through comprehensive validation, our method shows superior performance and the generated GuidingNet helps us to interpret DNMTs’ binding mechanism in different cell types and different species.

Our approach is outstanding for predicting DNMTs’ binding in the following aspects. First, GuidingNet integrates TF protein-protein network and co-expression across tissue and cell types to generate the weight of features in an unsupervised manner. This is different with existing procedures to select weights by computing an initial estimate using the response variable. Second, our method has a strong ability to select features by 1) setting cutoff in the GuidingNet and 2) fitting the regularized logistic regression. Third, we provide further biological insights into the difference of binding mechanism of DNMTs in different cell types. Fourth, our method is capable of making across-tissue predictions and validation by independent WGBS data. Last, GuidingNet is general enough to predict other chromatin regulators’ binding both in the human and mouse and shows outstanding performance.

Our model can further be improved from many aspects. First, the missing of gold-standard data limits us from training models in more tissues and cell types. Second, several studies have demonstrated that DNMT3 binding is closely related to chromatin histone modification, such as DNMT3B preferentially targets gene bodies marked with H3K36me3 [19, 20]. Many of the ChIP-seq of histone modification are available. Integrating other histone modification information in our model may benefit the prediction accuracy. Third, histone methyltransferases and demethylase also play an important role in mammal, such as KMT and KDM family are associated with embryo development and cancer occurrence and progression [21, 22]. Last, alterations in DNA methylation patterns have been implicated in embryo development and tumorigenesis in many studies [23-25]. We look forward to deciphering the regulatory mechanism of mediating DNA methylation status alteration and how it impacts these important biological processes.

## Acknowledgements

This study was supported by the National Science Foundation of China Grants 11871463, 61671444, and 61621003, Shanghai Municipal Science and Technology Major Project (No. 2017SHZDZX01), CAS “Light of West China” Program (No.xbzg-zdsys-201913), the “CAS Interdisciplinary Innovation Team” project.. CG is supported by Research Program of science and technology at Universities of Inner Mongolia Autonomous Region No NJZY19005.

## Author Contributions

Conceptualization: Yong Wang.

Investigation: Lixin Ren, Caixia Gao, Zhana Duren

Methodology: Lixin Ren, Caixia Gao, Zhana Duren, Yong Wang.

Software: Lixin Ren

Writing: Lixin Ren, Caixia Gao, Zhana Duren, and Yong Wang.

## Supporting information

**S1 Fig. The physical process that transcription factors and co-factor recruits DNMTs to a particular locus to methylate cytosines**.

One of the possible mechanisms is that transcription factors recognize specific DNA motifs in a genomic region and recruit DNMTs to methylate cytosines in that region. Black and white circles represent methylated and unmethylated sites, respectively.

**S2 Fig. The performance of ordinary logistic regression by using all three features is similar to GuidingNet**.

ROC curves of ordinary logistic regression prediction binding by using all three features for DNMT3A in human MCF-7 cells, DNMT3B in human HepG2 cells, DNMT3B in human foreskin keratinocyte, and Dnmt3b and Dnmt3l in E14 mESC.

**S3 Fig. Comparison of the performance between GuidingNet and randomly selected TFs for prediction**.

(A-E) The boxplot shows the AUC values of 1000 times randomly selected TFs for prediction in five scenarios and the black line is the AUC value of our method.

**S4 Fig. The performance of a single feature for predicting 4 CRs in the mouse E14**.

ROC curves of a single feature and all three features (openness, GC content, and TF information) prediction of binding for Ep300, Ezh2, Setdb1, and Suz12.

**S5 Fig. Overview of protein-protein interaction networks, the candidate GuidingNet, and the GuidingNet of 5 Ep300 in mouse E14**.

(A-C) The protein-protein interaction networks, the candidate GuidingNet, and the GuidingNet of Ep300. The cutoff used to generate the candidate GuidingNet was 0.1. The blue nodes in protein-protein interaction networks indicate the CR. The red edges indicate the direct interaction between the CR with TFs. The width of the edge in the candidate GuidingNet indicates its weight. The wider the width of the edge, the greater its weight. (D) Bar plot indicates the number of TFs in the PPI network (PPI), the candidate GuidingNet (CGN), and the GuidingNet (GN).

**S6 Fig. Overview of protein-protein interaction networks, the candidate GuidingNet, and the GuidingNet of Ezh12 in mouse E14**.

(A-C) The protein-protein interaction networks, the candidate GuidingNet, and the GuidingNet of Ezh2. The cutoff used to generate the candidate GuidingNet was 0.1. The blue nodes in protein-protein interaction networks indicate the CR. The red edges indicate the direct interaction between the CR with TFs. The width of the edge in the candidate GuidingNet indicates its weight. The wider the width of the edge, the greater its weight. (D) Bar plot indicates the number of TFs in the PPI network (PPI), the candidate GuidingNet (CGN), and the GuidingNet (GN).

**S7 Fig. Overview of protein-protein interaction networks, the candidate GuidingNet, and the GuidingNet of Setdb1 in mouse E14**.

(A-C) The protein-protein interaction networks, the candidate GuidingNet, and the GuidingNet of Setdb1. The cutoff used to generate the candidate GuidingNet was 0.1. The blue nodes in protein-protein interaction networks indicate the CR. The red edges indicate the direct interaction between the CR with TFs. The width of the edge in the candidate GuidingNet indicates its weight. The wider the width of the edge, the greater its weight. (D) Bar plot indicates the number of TFs in the PPI network (PPI), the candidate GuidingNet (CGN), and the GuidingNet (GN).

**S8 Fig. Overview of protein-protein interaction networks, the candidate GuidingNet, and the GuidingNet of Suz12 in mouse E14**.

(A-B) The protein-protein interaction networks and the candidate GuidingNet of Suz12. The cutoff used to generate the candidate GuidingNet was 0.1. The blue nodes in protein-protein interaction networks indicate the CR. The red edges indicate the direct interaction between the CR with TFs. The width of the edge in the candidate GuidingNet indicates its weight. The wider the width of the edge, the greater its weight. (C) Bar plot indicates the number of TFs in the PPI network (PPI), the candidate GuidingNet (CGN), and the GuidingNet (GN).

**S9 Fig. The performance of single feature for predicting 6 CRs in human HepG2 cells**.

ROC curves of a single feature and all three features (openness, GC content, and TF information) prediction of binding for EP300, EZH2, HDAC2, SMARCC2, SMARCE1, and SUZ12.

**S10 Fig. Overview of protein-protein interaction networks, the candidate GuidingNet, and the GuidingNet of EP300 in human HepG2**.

(A-C) The protein-protein interaction networks, the candidate GuidingNet, and the GuidingNet of a CR. The cutoff used to generate the candidate GuidingNet was 0.5. The blue nodes in protein-protein interaction networks indicate the CR. The red edges indicate the direct interaction between the CR with TFs. The width of the edge in the candidate GuidingNet indicates its weight. The wider the width of the edge, the greater its weight. Black, blue and red bars indicate the number of TFs in the PPI network (black), the candidate GuidingNet (blue), and the GuidingNet (red).

**S11 Fig. Overview of protein-protein interaction networks, the candidate GuidingNet, and the GuidingNet of EZH2 in human HepG2**.

(A-C) The protein-protein interaction networks, the candidate GuidingNet, and the GuidingNet of EZH2. The cutoff used to generate the candidate GuidingNet was 0.5. The blue nodes in protein-protein interaction networks indicate the CR. The red edges indicate the direct interaction between the CR with TFs. The width of the edge in the candidate GuidingNet indicates its weight. The wider the width of the edge, the greater its weight. Black, blue and red bars indicate the number of TFs in the PPI network (black), the candidate GuidingNet (blue), and the GuidingNet (red).

**S12 Fig. Overview of protein-protein interaction networks, the candidate GuidingNet, and the GuidingNet of HDAC2 in human HepG2**.

(A-C) The protein-protein interaction networks, the candidate GuidingNet, and the GuidingNet of HDAC2. The cutoff used to generate the candidate GuidingNet was 0.5. The blue nodes in protein-protein interaction networks indicate the CR. The red edges indicate the direct interaction between the CR with TFs. The width of the edge in the candidate GuidingNet indicates its weight. The wider the width of the edge, the greater its weight. Black, blue and red bars indicate the number of TFs in the PPI network (black), the candidate GuidingNet (blue), and the GuidingNet (red).

**S13 Fig. Overview of protein-protein interaction networks, the candidate GuidingNet, and the GuidingNet of SMARCC2 in human HepG2**.

(A-C) The protein-protein interaction networks, the candidate GuidingNet, and the GuidingNet of SMARCC2. The cutoff used to generate the candidate GuidingNet was 0.5. The blue nodes in protein-protein interaction networks indicate the CR. The red edges indicate the direct interaction between the CR with TFs. The width of the edge in the candidate GuidingNet indicates its weight. The wider the width of the edge, the greater its weight. Black, blue and red bars indicate the number of TFs in the PPI network (black), the candidate GuidingNet (blue), and the GuidingNet (red).

**S14 Fig. Overview of protein-protein interaction networks, the candidate GuidingNet, and the GuidingNet of SMARCE1 in human HepG2**.

(A-C) The protein-protein interaction networks, the candidate GuidingNet, and the GuidingNet of SMARCE1. The cutoff used to generate the candidate GuidingNet was 0.5. The blue nodes in protein-protein interaction networks indicate the CR. The red edges indicate the direct interaction between the CR with TFs. The width of the edge in the candidate GuidingNet indicates its weight. The wider the width of the edge, the greater its weight. Black, blue and red bars indicate the number of TFs in the PPI network (black), the candidate GuidingNet (blue), and the GuidingNet (red).

**S15 Fig. Overview of protein-protein interaction networks, the candidate GuidingNet, and the GuidingNet of SUZ12 in human HepG2**.

(A-B) The protein-protein interaction networks and the candidate GuidingNet of Suz12. The cutoff used to generate the candidate GuidingNet was 0.1. The blue nodes in protein-protein interaction networks indicate the CR. The red edges indicate the direct interaction between the CR with TFs. The width of the edge in the candidate GuidingNet indicates its weight. The wider the width of the edge, the greater its weight. (C) Bar plot indicate the number of TFs in the PPI network (PPI), the candidate GuidingNet (CGN), and the GuidingNet (GN).

**S16 Fig. The better prediction of E14**.**5 is related to the right guiding TFs choice**.

(A) Venn diagram shows the number of different regions the between E13.5 and E14.5.

(B) Motif enriched Guiding TFs in the unique binding regions of E14.5. Red nodes indicate TFs have high gene expression in the liver of E14.5

**S17 Fig. Bar plot of the distance between predicted binding regions and the transcription start site (TSS) at 7 time points**.

**S1 Table. The description of RNA-seq data for mouse of our collection**.

**S2 Table. The description of RNA-seq data for human of our collection**.

**S3 Table. Literature validation of our guiding co-factors**.

**S4 Table. Comparison of GuidingNet with other alternative methods**.

